# Effects of Milk Fat Globule Epidermal Growth Factor VIII On Age-Associated Arterial Elastolysis, Fibrosis, and Calcification

**DOI:** 10.1101/2020.10.05.326728

**Authors:** Soo Hyuk Kim, Lijuan Liu, Leng Ni, Li Zhang, Jing Zhang, Yushi Wang, Kimberly R. McGraw, Robert Monticone, Richard Telljohann, Christopher H. Morrell, Edward G. Lakatta, Mingyi Wang

## Abstract

Milk fat globule-EGF factor 8 (MFG-E8) protein increases with age and is mainly secreted by vascular smooth muscle cells in the arterial wall. Here, we investigated the role of MFG-E8 signaling during proinflammation, elastolysis, fibrosis, and calcification within the aging arterial wall. In vivo studies indicated that (1) Elastic lamina breaks collagen deposition and calcium-phosphorus products were markedly increased in the aging arterial wall of rats; (2) MFG-E8 protein abundance was markedly increased while intact tropoelastin (TPELN), an element of repair of the elastic fibers, was markedly decreased in the aging arterial wall of rats; (3) The absence of MFG-E8 markedly alleviated age-associated increases in elastic lamina breaks, collagen deposition and calcium-phosphorus products in mice; and (4) MFG-E8 deficiency significantly decreased age-associated increases in matrix metalloproteinase type II (MMP-2) activation, alkaline phosphatase, and runt-related transcription factor 1 (Runx1) expression in the aortic walls of mice. The in vitro studies demonstrated that (1) treating either young or old rat VSMCs with recombinant human MFG-E8 protein (rhMFG-E8) significantly reduced TPELN levels while MFG-E8 gene silencing significantly increased TPELN levels; (2) rhMFG-E8 treatment activated MMP-2 levels in both young and old VSMCs; and (3) MMP-2 bound to and cleaved TPELN secreted from VSMCs. Thus, these findings suggest that MFG-E8 signaling promotes age-associated adverse structural remodeling, including elastolysis, fibrosis, and calcification; however, MFG-E8 deficiency markedly mitigates these adverse effects in mice.

## Introduction

Aging is the major risk factor for quintessential cardiovascular diseases, such as hypertension and atherosclerosis, mainly due to arterial proinflammatory remodeling.^1–3^ Proinflammation is sterile low-grade inflammation that accompanies aging, in which cytokines are secreted by nonprofessional vascular cells rather than professional immune cells.^1–3^ Arterial wall remodeling is associated with the phenotypic shift of vascular smooth muscle cells (VSMCs) from a contractile to synthetic proliferative, migratory, invasive, secretory, and senescent state, creating a heterogenous proinflammatory elastolytic, fibrotic, and calcified microenvironment.^1–12^ The arterial wall becomes less resilient and stiffened with advancing age due in part to a trilogy of interrelated remodeling events: elastolysis, fibrosis, and calcification.^2,13^

Tropoelastin (TPELN), a core protein of elastic fibers, is secreted mainly from VSMCs in the arterial wall. TPELN, fibril-rich microfibrils, and other accessory proteins are assembled into a complex elastic network on the surface of VSMCs.^14^ Elastolysis is the collapse of the elastic network within the aged arterial wall, which is due mainly to the enzymatic cleavage of TPELN and microfibrils. Importantly, TPELN is a monomeric soluble precursor of insoluble elastin and plays an important role in maintaining VSMCs contractile phenotype, elastic network integrity, transforming growth factor-beta 1(TGF-β1) fibrogenic signaling, and cardiovascular health.^4,7– 9,14,15^ TPELN secreted from VSMCs in the aortic wall is susceptible to be degraded by activated matrix metalloproteinase type 2 (MMP-2), which increases with advancing age.^15–18^ The breakdown of the elastic network is accompanied by a reduction of the levels of intact TPELN protein and release of elastin derived peptides, promoting fibrosis and calcification.^4,7,8,15,17,19^

VSMCs secrete milk fat <u>g</u>lobule epidermal <u>g</u>rowth factor VIII (MFG-E8), an extracellular multifunctional glycoprotein, also known as lactadherin in the arterial wall.^20,21^ MFG-E8 protein abundance is increased in the aging arterial wall and promotes adverse arterial wall remodeling through the processes of the inflammation, proliferation, and invasion of VSMCs.^20–24^ However, how effects of MFG-E8 on adverse extracellular matrix (ECM) remodeling, including elastolysis, fibrosis, and calcification, remains unknown.

In the present, in vivo studies indicate that MFG-E8 abundance is closely associated with the age-associated aortic wall elastic fiber degradation, calcification, and fibrosis in FXBN rats and wild type (WT) mice. Importantly, the absence of MFG-E8 alleviated the elastin breakdown, fibrosis, and calcification in the aging arterial wall of MFG-E8 KO mice. In the current in vitro studies demonstrate that MFG-E8 activates MMP-2 and subsequently cleaves TPELN secreted from VSMCs. These findings suggest that MFG-E8 signaling promotes age-associated adverse remodeling; however, the absence of MFG-E8 markedly retards age-associated adverse molecular, structural remodeling in mice.

## Materials & Methods

All data, analytic methods, and study materials will be made available to other researchers for purposes of reproducing the results or replicating the procedure. The material that supports the findings of this study are available from the corresponding author upon reasonable request.

## Experimental Animals

### Rats

All procedures were performed per protocol (413-LCS-2021) and approved by the National Institute on Aging (NIA) in accordance with the National Institutes of Health (NIH) Animal Care and Use Committee. Eight-month old (*n* = 16, young) and 30-month old (*n* = 16, old) male Fisher 344 crossbred Brown Norway (FXBN) rats were obtained from the NIA Aged Rodent Colonies. Animals were sacrificed and thoracic aortae were harvested as previously described.^25^ Fresh aortic tissue was snap-frozen in liquid nitrogen (LN_2_) for Western blotting analysis.

In addition, aortic tissues harvested from young and old FXBN rats (*n* = 12/age group) were flatly embedded in paraffin. Longitudinal or cross-sectional (5μm in thickness) aortic walls were utilized for histology and histochemistry staining, including hematoxylin & eosin (HE), Masson’s trichrome, elastic tissue fibers Verhoeff’s Van Gieson (EVG), Alizarin Red S, and von Kossa staining, according to manufacturer’s instruction (Polysciences, Inc. Warrington PA), and morphometric analyses (MetaMorph Microscopy Automation and Image Analysis Software, Molecular Devices, LLC, CA) as described previously.^17,25,26^

### Mice

All experiments were performed according to the protocol (<u>445-LCS-2022</u>) and approved by the NIA in accordance with the NIH Animal Care and Use Committee. MFG-E8 KO mice were obtained from Dr. Mark Udey at National Cancer Institute, and generated, characterized, and genotyped as described previously.^27,28^ Animals were housed and used in experiments in accordance with the NIH guidelines.

Animals were sacrificed and thoracic aortae were harvested for histopathology and protein analyses as previously described^25^ Thoracic aortic tissues were harvested from young MFG-E8 KO and age-matched WT (*n* = 12/genotype group, 40-week-old) and old MFG-E8 KO and age-matched WT animals (*n* = 12/genotype group, 96-week-old) were embedded in paraffin and sectioned for histology and histochemistry staining, including hematoxylin eosin for general overview, Masson’s trichrome for collagen, EVG for elastin fibers, Alizarin Red S for calcium deposits, according to manufacturer’s instruction (Polysciences, Inc. Warrington, PA, USA), and morphometric analyses described previously.^17,25,26^ Fresh aortic tissue was harvested from MFG-E8 KO and WT animals was snap-frozen in liquid nitrogen (LN_2_) for Western blotting analysis.

### Primary VSMCs Isolation and Culture

All in vitro experiments were performed on cultured rat VSMCs which were enzymatically isolated from aortae of rats aged 8 months (young) and 30 months (old), as previously described.^18,23^ Briefly, FXBN rat thoracic aortae were rinsed in Hanks balanced salt solution (HBSS) containing 50 μg/mL penicillin, 50 μg/mL streptomycin, and 0.25 μg/mL amphotericin B (Thermo Fisher Scientific, Lanham, MD, USA). After digestion for 30 min in 2mg/mL of collagenase I solution (Worthington Biomedical, Freehold, NJ, USA) at 37°C, the adventitia and intima were removed from the vessel medial layer and then were further digested with a mix of 2mg/mL collagenase II/0.5mg/mL elastase (Sigma, Burlington, MA, USA) for 90–120 min at 37°C. Isolated VSMCs were washed and plated in complete medium containing 20% fetal bovine serum (FBS) and cultured at 37°C. Cells were sub-cultured (passage 3–5) in modified DMEM with 10% FBS. After serum removal for 24 hours, cells cultured in 2.5% FB were treated with recombinant human MFG-E8 (rh-MFG-E8, R&D Systems, Inc., Minneapolis, MN, USA).

### Western Blotting

For Western blotting, total 10 μg of whole cell or arterial protein lysates were resolved by SDS-PAGE and transferred to a nitrocellulose membrane (Bio-rad, Hercules, California, USA). The transferred membranes were incubated in phosphate-buffered saline (PBS) containing primary antibodies at 4°C for 24 hours. Horseradish peroxidase (HRP)-conjugated IgG (Abcam, Cambridge, MA, USA) was used as a secondary antibody and detected with SuperSignal West Pico Chemiluminescent Substrate (Pierce Biotechnology, Rockford, IL, USA). Densitometric density of the bands was analyzed by ImageJ. Primary antibodies are listed in **Table 1**.

**Table 1.** Primary Antibodies.

| Antibody | Species | Titer blotting | Titer staining | Cat# | Source |
| --- | --- | --- | --- | --- | --- |
| β-actin | M | 1:1000 | | sc-47778 | Santa Cruz, CA |
| MFG-E8 | R | 1:200 |  | sc-33546 | Santa Cruz, CA |
| MMP-2 | R | 1:1400 |  | NB200-114 | Novus Bio, CO |
| Tropoelastin | R | 1:500 |  | orb13391 | Biorbyt, UK |
| Tropoelastin | R |  | 1:50 | ab21600 | Abcam, MA |

M = mouse, R = rat

### Quantitative Real-Time PCR

Messenger ribonucleic acid (mRNA) was extracted from the aortic tissue or from early passage primary VSMCs in culture isolated from young and old rat aortae using the RNA precipitation kit (Qiagen, Germantown, MD, USA). RNA (200ng) was isolated according to the manufacturer’s instructions (Applied Biosystems, Foster City, CA, USA). All the primers used for Real-Time PCR (RT-PCR) analysis was designed using Primer Express software 2.0 (Applied Biosystems, Foster City, CA, USA), and synthesized by Thermo Fisher Scientific (Carlsbad, CA, USA). The following primer sequences were used: TPELN, forward ATCGGAGGTCCAGGCATTG; and backward ACCAGCACCAACCCCGTAT. MFG-E8, forward ACACACAGCGAGGGGACAT; and backward ATCTGTGAATCGGCAATGG. H1f0, forward AGCCACTACAAGGTGGGTGAGA; and backward TTGAGAACACCGGTGGTCACT. Real-time PCR was performed according to the SYBR Green PCR protocol (Applied Biosystems, Foster City, CA, USA): 10 min at 95°C (one cycle); 30 second (sec) at 95°C; 30 sec at 60°C; and 30 sec at 72°C (40 cycles). Gene-specific PCR products were continuously measured by an ABI PRISM 7900 HT Sequence Detection System (PE Applied Biosystem Norwalk, CT, USA). The PCR product sizes were verified by agarose gel electrophoresis. Samples were normalized to the expression of the “housekeeping” gene, H1F0.

### Transfection of Small-Interfering RNA

Small RNA silencing MFG-E8 and MMP-2 were synthesized by Ambion of Thermo Fisher Scientific (Lanham, MD, USA). The following primer sequences were used: MFG-E8 forward GAACAUCUUUGAGAAACCUTT; and backward AGGUUUCUCAAAGAUGUUCTT. MMP-2 forward CCACUACGCUUUUCUCGAATT; and backward UUCGAGAAAAGCGUAGUGGAG. VSMCs were transfected for 48 hours with scrambled siRNA (20 nM) or targeted protein siRNA (20 nM), using liposome based RNAiMAX (Thermo Fisher Scientific, Lanham, MD, USA), according to the manufacturer’s instructions.

### In Situ Gelatin Zymography

VSMCs in situ zymography were performed according to a modified protocol, as described previously.^29^ In brief, VSMCs were isolated from young and old rat aortae. Cells were primarily cultured on a sterile coverslip after earlier three passages. Nuclear dye DAPI was added to a mixture of an equal volume of 0.5 μg/ml FITC-gelatin and agarose solution in a reactive buffer, including 1% Triton X-100, 50 mM Tris-HCl, 5 mM CaCl_2,_ and 1 μM ZnCl_2_. The early passage VSMCs covered with the mixed DAPI-FITC-gelatin solution were cultured and incubated in the dark at 37°C for 6 hours. The nuclei of VSMCs were stained with 4’,6-diamidino-2-phenylindole (DAPI, blue color). Notably, under fluorescence microscope, the FITC-gelatin substrate was cleaved by activated gelatinases appearing as green color.

### PAGE Gelatin Zymography

To detect MMP-2 activity, as secreted by VSMCs, equal amounts of homogenous protein from VSMCs (10 μg) or aortic walls (10 μg) and 2x zymogram sample buffer were mixed and loaded onto zymography gels (Invitrogen Novex Zymogram gels, Thermo Fisher Scientific, Lanham, MD, USA). The gels were renatured by incubation with Novex renaturing buffer for 30 minutes at room temperature and incubated in Novex developing buffer at 37°C overnight. Finally, the gel was stained with 0.5% Coomassie blue, and then de-staining to visualize the proteolytic lysis bands, which were analyzed using ImageJ software.

### Statistical Analysis

All data were presented as both individual values and mean ± SEM. Statistical analyses used two-way or two-way repeated measure ANOVA followed by Turkey post hoc test used for multiple comparisons. These statistical analyses were performed using GraphPad Prism (version 9.0) software. A value of *p* ≤ 0.05 was considered statistically significant.

## Results

### MFG-E8 and Adverse Aortic Remodeling with Aging

With advancing age, aortic wall was adversely structurally functionally remodeled, including elastic fiber degeneration, collagen deposition, and calcification, contributing to the increases of the morbidity and mortality of cardiovascular disease and shortening the lifespan.^1–3^ The current in vivo studies demonstrated that MFG-E8 abundance was closely associated with the arterial adverse remodeling in either rats or WT mice with aging (**Figures 1–7**). Importantly, MFG-E8 deficiency significantly alleviated the age-associated adverse remodeling in KO mice (**Figures 4– 7**).

**Figure 1.**
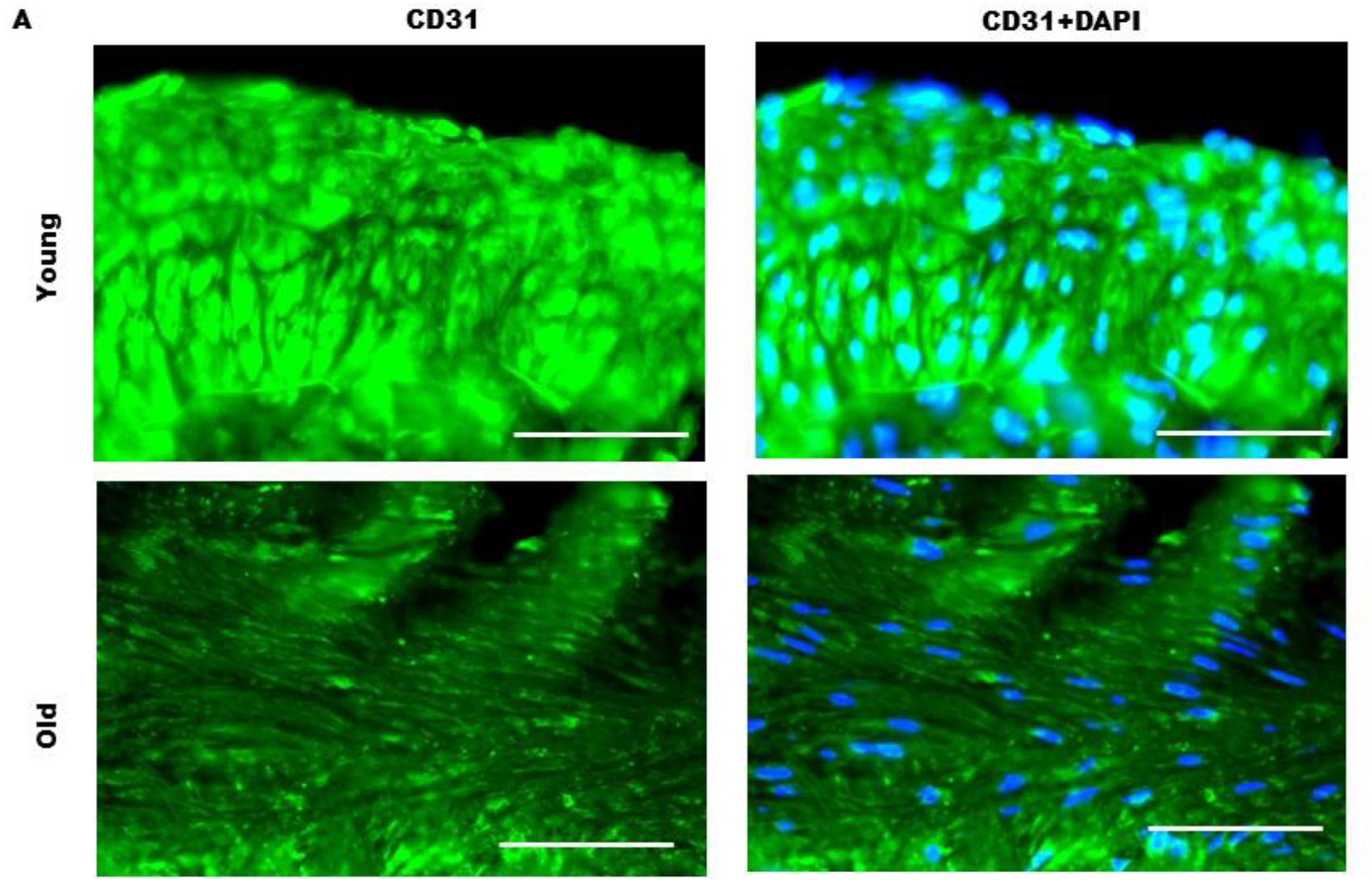
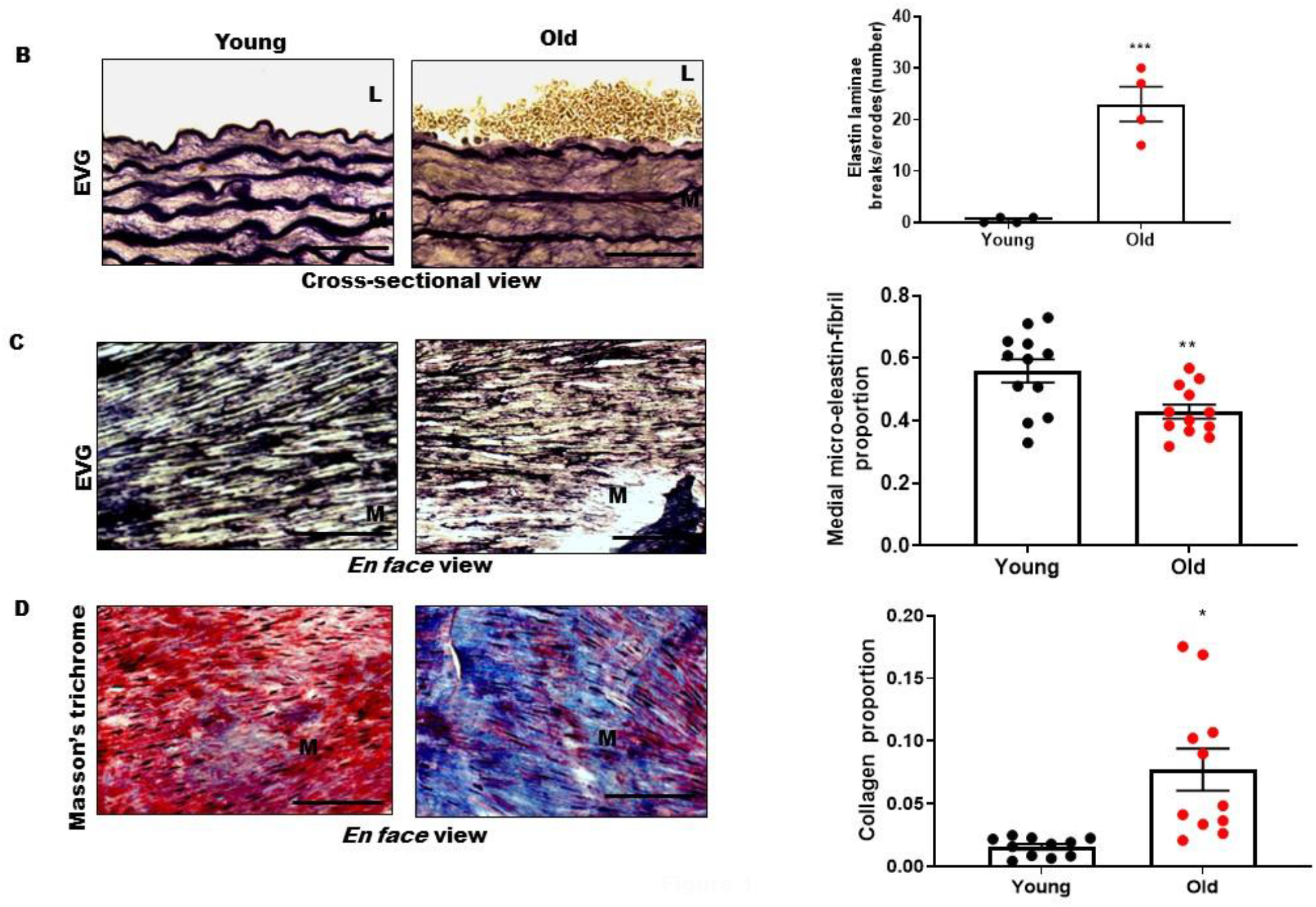
Elastolysis and fibrosis increases in aging rat aortic walls. **A**. Representative photomicrographs (left side) of en face views of rat aortic endothelium stain with CD31 (green) and DAPI (blue). **B**. Representative photomicrographs (left side) of cross-sectional views of rat aortic walls stained with EVG (dark color). Graph showing that individual numbers of aortic elastic laminae break/cross-section and mean ± SEM. \**p* < 0.05 by unpaired *t* test. **C**. Representative photomicrographs (left side) of en face view of aortic medial walls stained using EVG. Graph showing that individual values of elastic fiber proportion under high power view and mean± SEM. \**p* < 0.05 by unpaired *t* test. **D**. Representative photomicrographs (left side) of en face view of aortic medial wall stained with Masson’s trichrome (muscle, red color; extracellular matrix-collagen, blue color). Graph showing that individual values of collagen proportion under high power view and mean ± SEM. \**p* < 0.05 by unpaired *t* test. L, lumen; M, medium. Bar = 100μm.

#### Elastolysis, Fibrosis, and Calcification Increase in the Aging Rat Aortic Wall

Aging remodels the arterial wall. To determine the integrity of endothelium, immunofluorescence staining CD31 was performed (**Figure 1A)**, an en face view indicated that endothelial layer was defect (lower or absence of green color) in old vs young aortic wall (**Figure 1A**). To study the extent of elastolysis, we performed morphometrical analysis of histochemical EVG staining of cross-sectional and en face rat aortic walls. We observed that the breaks in the elastic meshwork of aortic elastin laminae, an index of elastolysis, was dramatically increased in old vs young rats (**Figure 1B)** while the inter-lamina micro-elastic fibers proportion of the elastic fraction, another index of elastolysis, was substantially decreased (**Figure 1C**). In addition, there was a significant increase of collagen deposition in the aortic wall in old vs young rats (**Figure 1D)** as determined by Masson’s trichrome staining.

Notably, other important evidence of calcification was made with Alizarin red staining, that demonstrated that calcium deposits were substantially elevated in old vs young aortic walls (**Figure 2A**); and Von Kossa staining demonstrated that phosphorus deposits were significantly elevated in old vs young aortic walls (**Figure 2B**).

**Figure 2.**
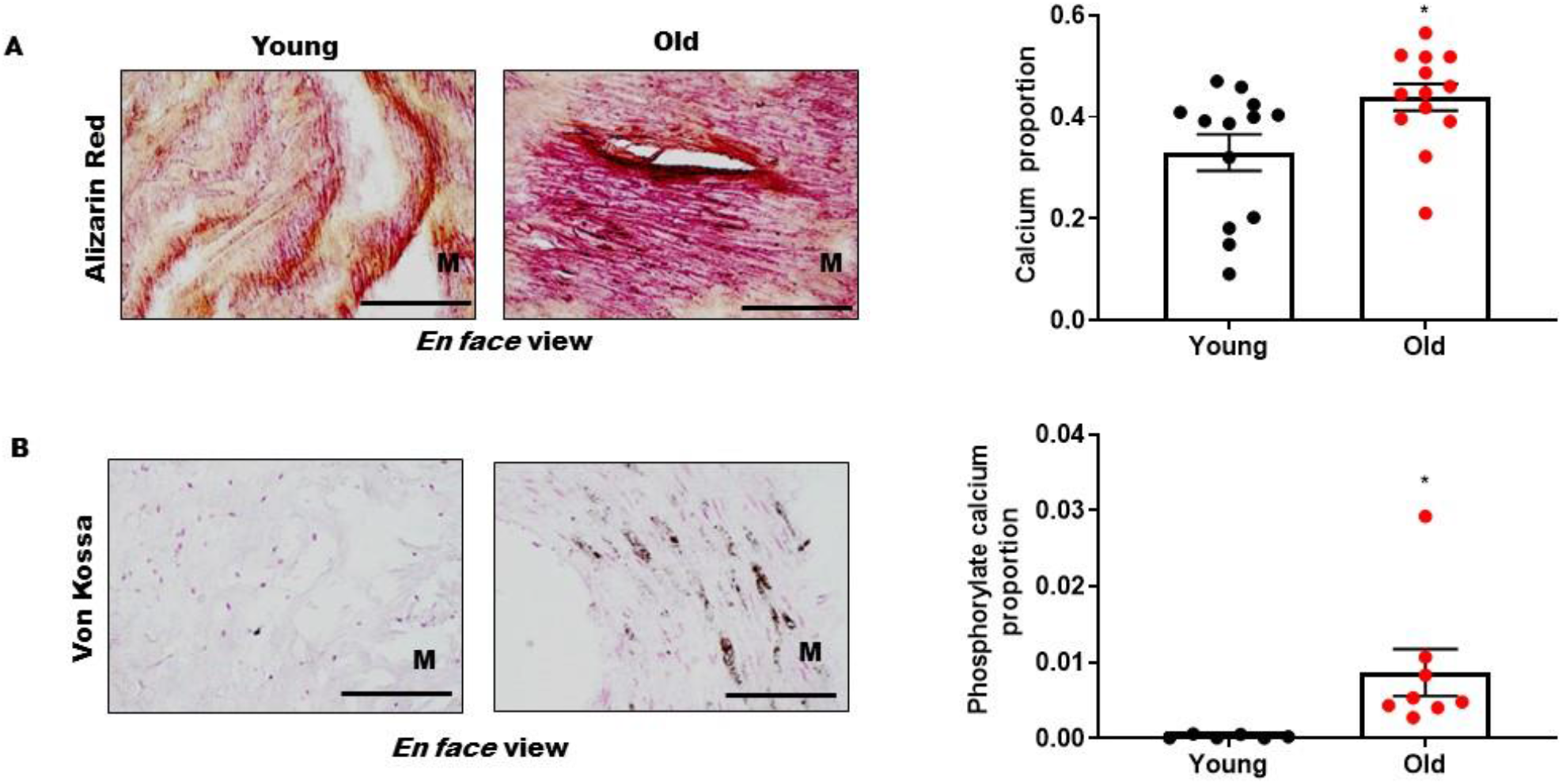
Calcification increases in aging rat aortic walls. **A**. Representative photomicrographs (left side) of en face view of rat aortic medial wall stained with Alizarin red (calcium deposits, red color). Graph showing that individual values of calcium deposit proportion under high power view and mean ± SEM. \**p* < 0.05 by unpaired *t* test. **B**. Representative photomicrographs (left side) of en face view of aortic medial wall stained with Von Kossa (phosphorylate calcium deposits, dark color). Graph showing that individual values of phosphorylate calcium deposit proportion and mean ± SEM. \**p* < 0.05 by unpaired *t* test. L, lumen; M, medium. Bar = 100μm.

#### Altered MFG-E8 and Tropoelastin Expression in the Aging Rat Aortic Wall

MFG-E8 and TPELN are inversely altered in the proinflamed aging arterial wall. Immunoblotting analyses showed that intact TPELN protein markedly decreased while MFG-E8 levels markedly increased in old vs young aortic walls (**Figure 3A & 3B)**. Immunofluorescence staining showed that MFG-E8 protein (red) levels increased while TPELN (green) protein levels were reduced in old vs young aortic walls (**Figure 3C, left panels**). Further, dual immunolabelling and morphometrical analysis showed that there was also a marked increase in the colocalization of MFG-E8 and TPELN in old vs young aortic walls (**Figure 3C, right panel**).

**Figure 3.**
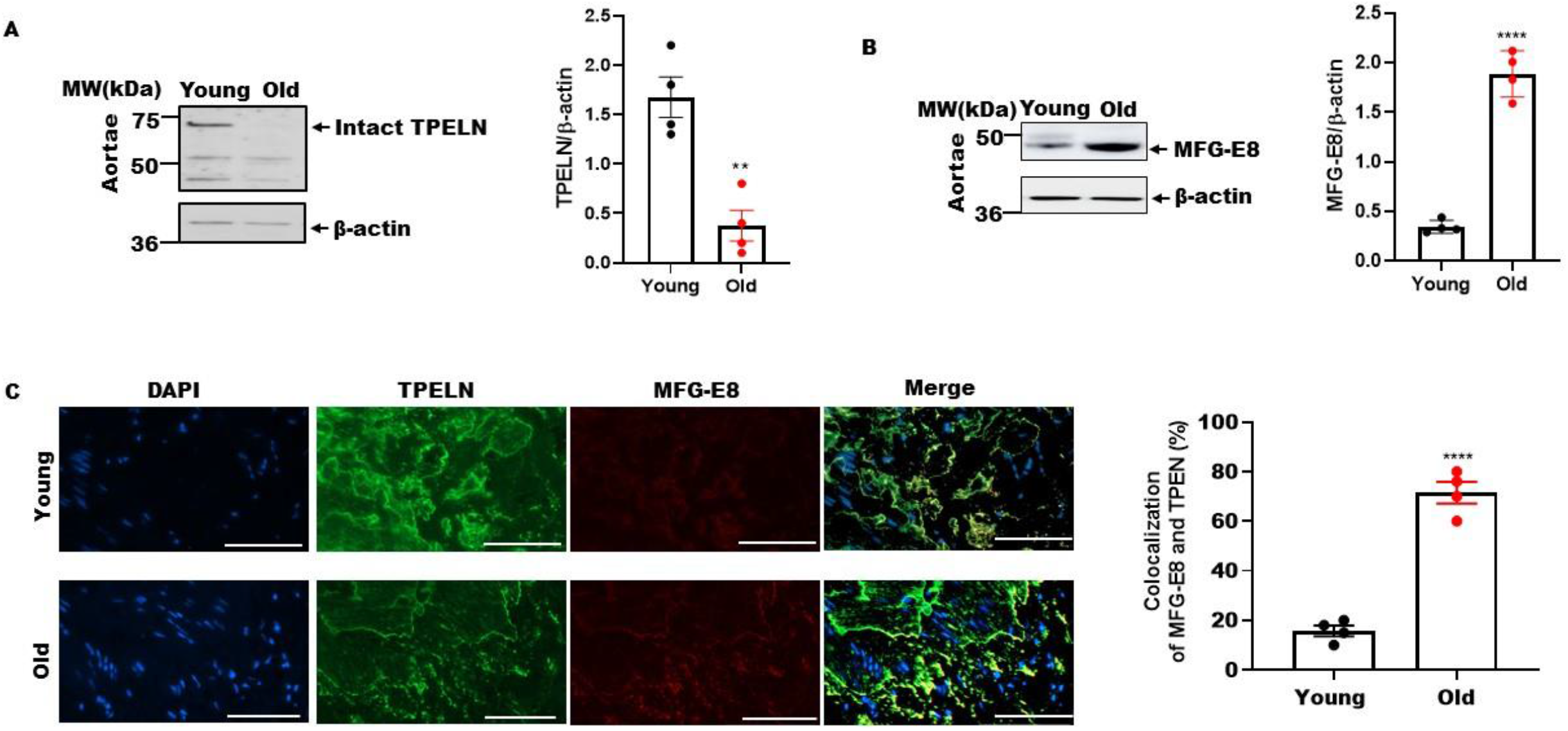
MFG-E8 and TPELN expression levels in aging aortae. **A**. Representative Western blot images of intact TPELN in young and old rat aortae. Graph showing that individual values of relative TPELN abundance and mean ± SEM. **B**. Representative Western blot images of MFG-E8 in young and old rat aortae. Graph showing that individual values of relative MFG-E8 protein expression and mean± SEM. **C**. Representative photomicrographs of en face view of aortic medial wall stained with TPELN (green color) and MFG-E8 (red color) as well as merge (yellow color) in young (upper panels) old (lower panels). Nuclei were counter-stained with DAPI (blue color). Bar = 100 μm. Graph showing that individual values of the percentage of colocalization of MFG-E8 and TPELN and mean ± SEM. \*\**p* < 0.001 by unpaired *t* test.

#### MFG-E8 Deficiency Impedes Adverse Structural Remodeling in Aging Mouse Aortic Walls

Our results indicated that MFG-E8 is a potential initiator of adverse arterial wall remodeling, i.e., elastic fiber degradation, we reasoned that adverse arterial wall remodeling would be substantially reduced in an environment where MFG-E8 is suppressed. To this end, we compared elastin degradation, collagen, and calcification in the arterial wall young (40 weeks) and old (96 weeks) WT and age-matched MFG-E8 KO mice.

Western blot analysis indicated that MFG-E8 protein was deficient in the aortic walls of homogeneous MFG-E8 KO mice but was enriched and significantly increased in the aortic walls of WT mice with aging (two-way ANOVA: main effect of age, *p* < 0.0001; main effect of genotype, *p* < 0.0001; and interaction of age and genotype, *p* < 0.0001) **(Figure 4A & 4B)**. Importantly, Western blot analysis indicated that aortic intact TPELN abundance was markedly decreased in old compared with young WT mice; however, the absence of MFG-E8 significantly alleviated this age effect (two-way ANOVA: main effect of age, *p* < 0.001; main effect of genotype, *p* < 0.05; and interaction of age and genotype, *p* < 0.05) (**Figure 4A & 4B)**. Interestingly, aortic intact elastin proportion (another index of elastin degradation) significantly decreased in old versus young in WT; however, MFG-E8 deficiency significantly lessens this age effect (two-way ANOVA: main effect of age, *p* < 0.0001; main effect of genotype, *p* < 0.0001, and interaction of age and genotype, *p* > 0.05) (**Figure 4C**).

**Figure 4.**
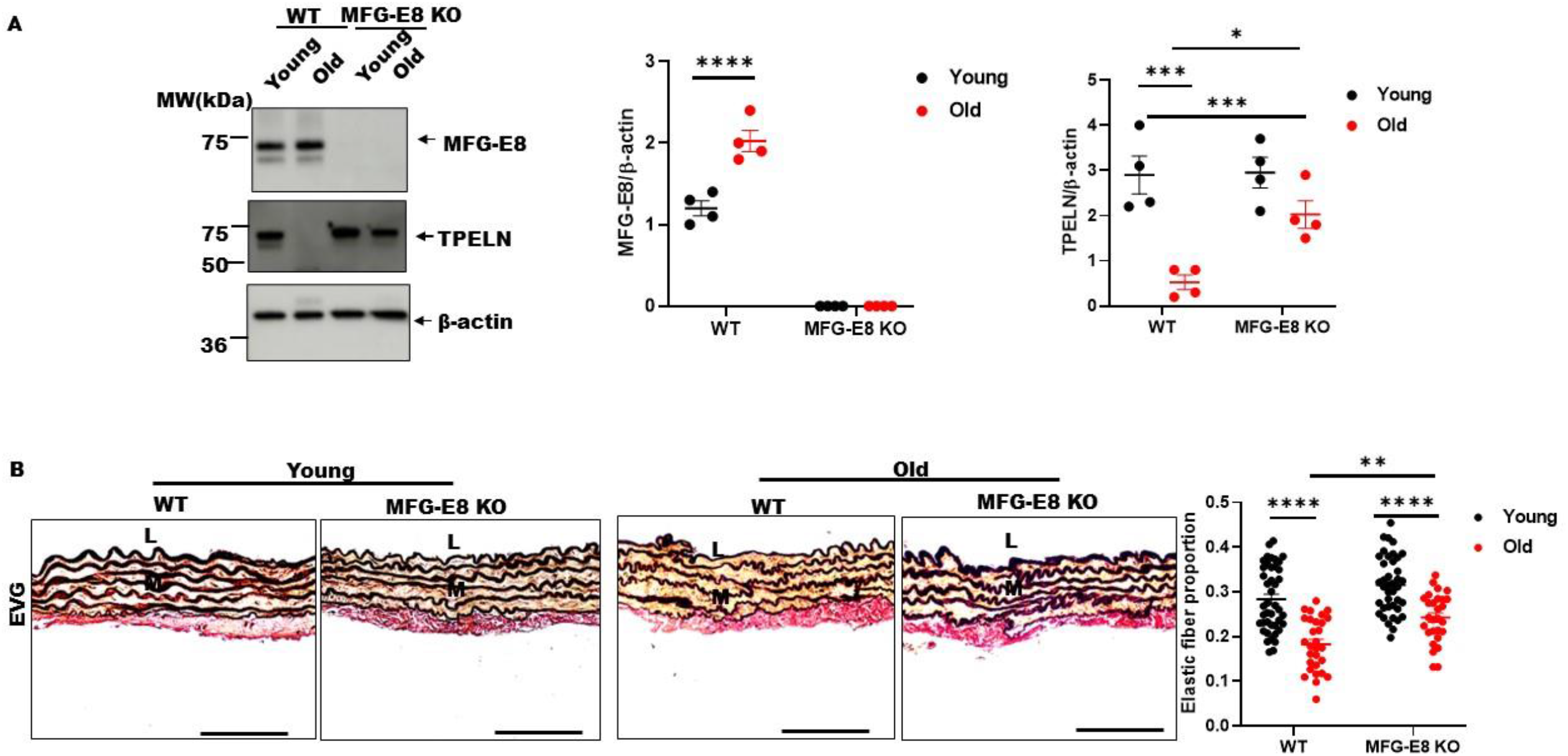
MFG-E8 and TPELN expression and elastic fiber degradation in aging mice aortic walls. **A**. Western blots of aortic MFG-E8 and TPELN proteins in young (40 weeks) and old (96 weeks) WT and MFG-E8 KO mice. Graph (middle panel) showing that individual values of relative MFG-E8 expression and mean± SEM, analyzed with two-way ANOVA (*p* < 0.0001, main effect of age; *p* < 0.001, main effect of genotype; *p* < 0.001 interaction of age x genotype). Graph (right panel) upper showing that individual values of relative TPELN expression and mean ± SEM, analyzed with two-way ANOVA (main effect of age, *p* < 0.001; main effect of genotype, *p* < 0.5; interaction of age *x* genotype, *p* < 0.05). **B**. Representative photomicrographs of cross-sectional view of aortic walls stained using EVG (dark color). Graph showing that individual values of elastin proportion/high power field and mean ± SEM, analyzed with two-way ANOVA: age, *p* < 0.0001, genotype, *p* < 0.0001; no interaction of age and genotype interaction, *p* > 0.05. Post hoc Turkey multiple comparison test: \**p* < 0.05 and \*\**p* < 0.01.

In contrast, age increased collagen proportion in WT animals while MFG-E8 KO abolished this age effect (main effect of age: *p* < 0.05; main effect of genotype: *p* > 0.05; interaction of age and genotype: *p* < 0.01) (**Figure 5A)**. Importantly, age increased calcium deposits proportion in WT animals, but MFG-E8 KO significantly alleviates this age effect (main effect of age: *p* < 0.0001; main effect of genotype: *p* > 0.01; interaction of age and genotype: *p* < 0.05) (**Figure 5B)**.

**Figure 5.**
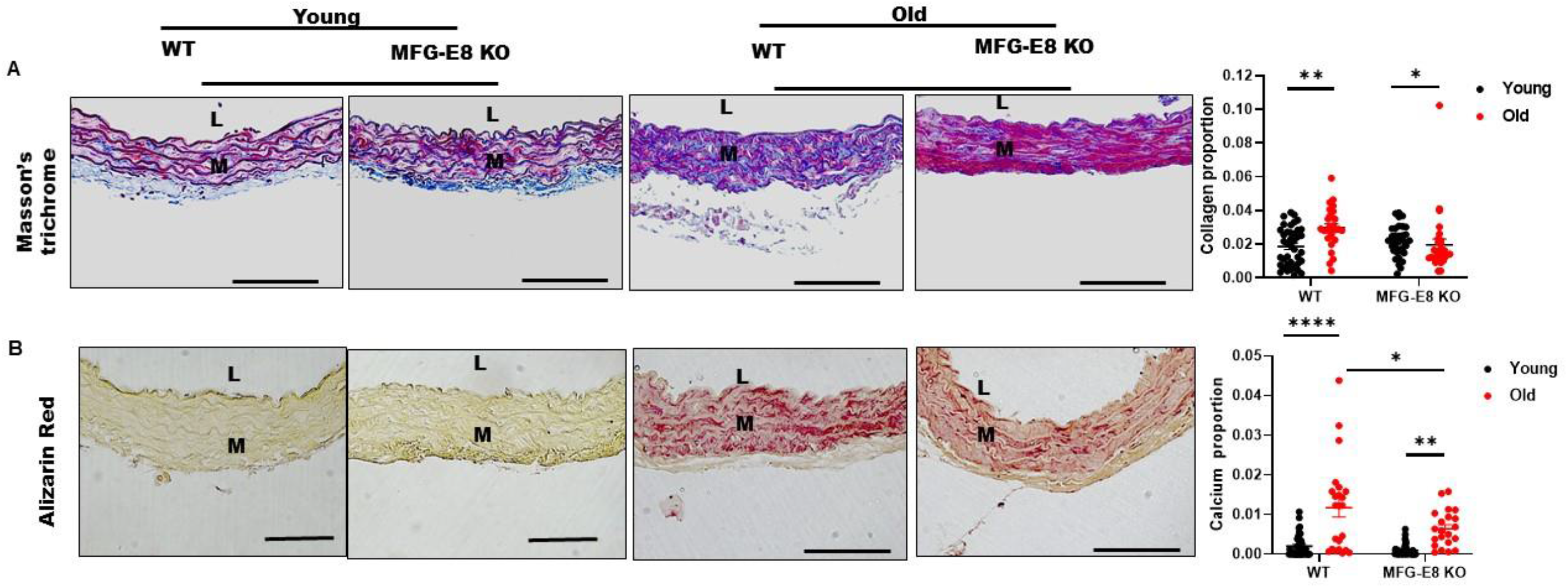
Adverse aortic wall collagen deposition and calcification in aging mice. **A**. Representative photomicrographs of cross-sectional view of aortic medial walls stained with Masson’s trichrome from young and old MFG-E8 KO and WT mice (muscle, red color; extracellular matrix, blue color). Graph showing that individual values of collagen proportion/high power field and mean ± SEM, analyzed with two-way ANOVA: age, *p* < 0.05; age and genotype interaction, *p* < 0.01. Post hoc Turkeys multiple comparison test: \**p* < 0.05 and \*\**p* < 0.01. **B**. Representative photomicrographs of cross-sectional view of aortic medial walls stained with Alizarin red (calcium deposits, red color). Graph showing that individual values of calcium deposition and mean± SEM analyzed with two-way ANOVA: age, *p* < 0.001; age and genotype interaction, *p* < 0.05. Post hoc Turkey multiple comparison test: \**p* < 0.05, \*\**p* < 0.01, and \*\*\*\**p* < 0.0001. L, lumen; M, medium. Bar = 100 μm.

#### MFG-E8 Deficiency Minimizes Calcifying Molecular Remodeling in Aged Mouse Aortic Walls

Activated MMP-2, alkaline phosphatase (ALP), and runt-related transcription factor 2 (RUNX2) are key molecules to the formation of calcified elastic fragments and osteochondrogenic transdifferentiated VSMC.^9,11,12,30,31^ Western blotting analysis of aortic proteins demonstrated that activated MMP-2 protein levels and activity were markedly increased in old WT vs old MFG-E8 KO mice (**Figure 6A and B**). The pro-calcification proteins ALP and RUNX2 were markedly increased in old WT versus MFG-E8 KO (**Figure 6C**).

**Figure 6.**
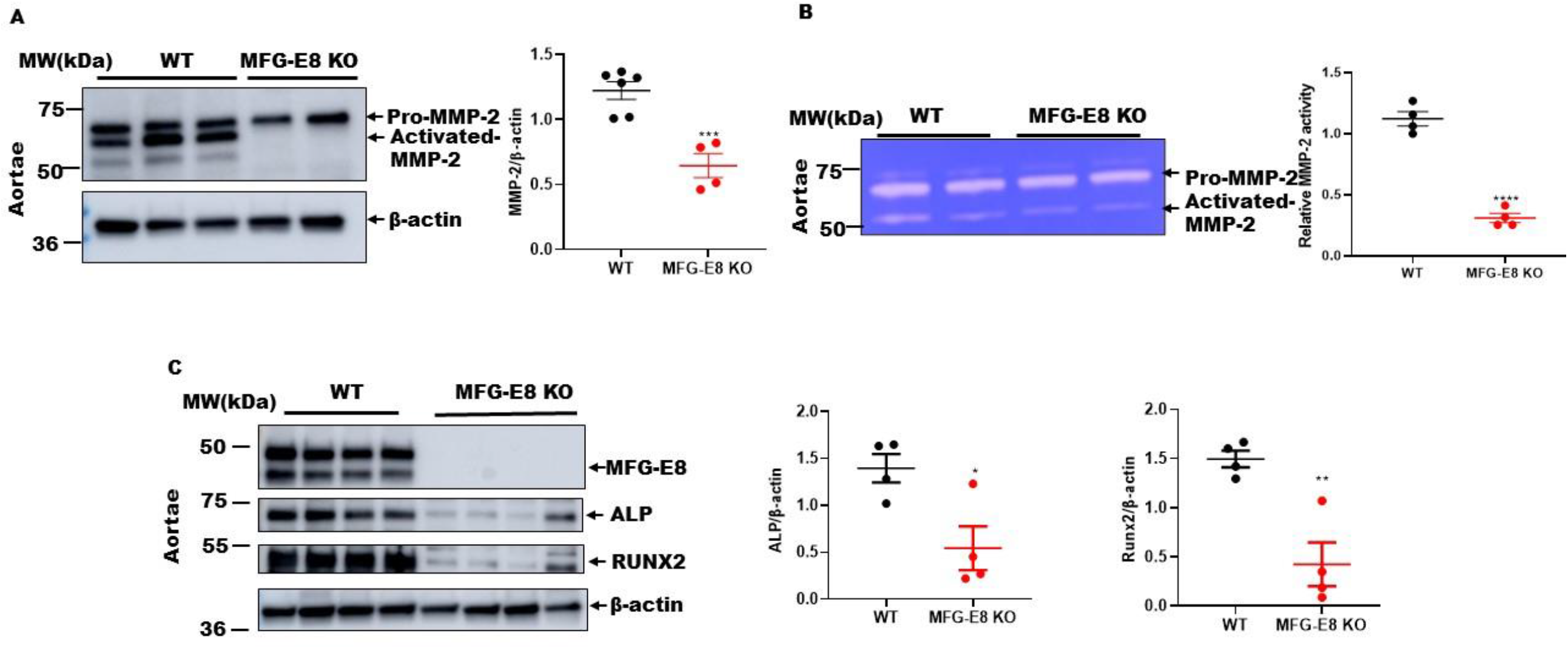
Arterial calcification molecules in aging mice. **A**. Representative Western blot of homogenous aortic protein from MFG-E8 KO and WT mice for MMP-2. Graph showing that individual values of relative MMP-2 abundance and mean ± SEM. Unpaired *t* test: \*\*\**p* < 0.001; \*\*\*\**p* < 0.0001. **B**. PAGE zymogram of homogenous aortic protein from MFG-E8 KO and WT mice for MMP-2. Graph showing that individual values of relative activated MMP-2 and mean ± SEM. Unpaired *t* test, \*\*\**p* < 0.001; \*\*\*\**p* < 0.0001 **C**. Western blots of homogenous aortic protein for ALP, and RUNX2 from old MFG-E8 KO and age-matched WT mice. Graph showing that individual relative ALP and RUNX2 values and mean ± SEM. Unpaired *t* test, \**p* < 0.05; \*\*\**p* < 0.01.

### MFG-E8 Mediates VSMC Proinflammation with Aging

VSMCs are predominant cells in the media of aortic wall. The current in vitro and ex vivo results indicated that age-associated increases in MFG-E8, MMP-2 activation, and elastin degradation in VSMCs play crucial roles in age-associated arterial remodeling **(Figures 7–10)**.

**Figure 7.**
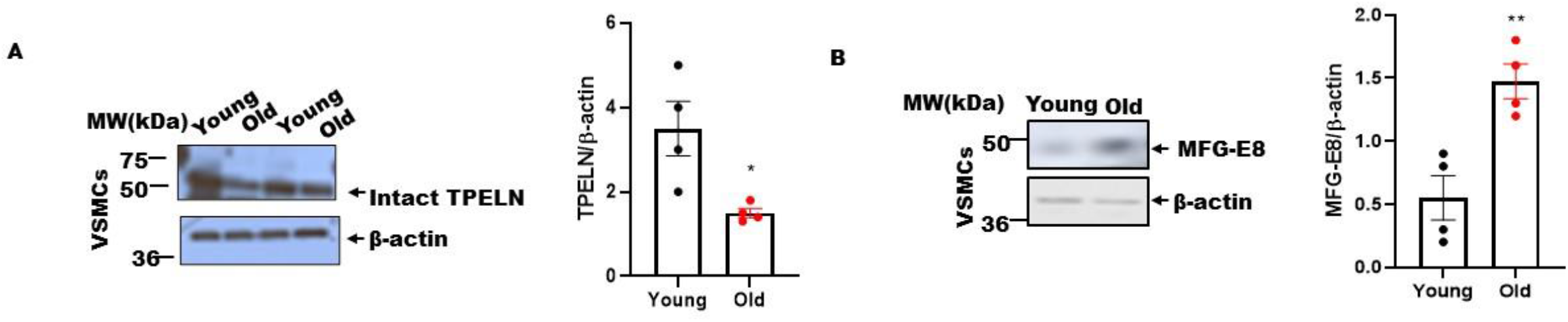
The age-associated changes in TPELN and MFG-E8 expression in VSMCs. **A**. Representatives Western blots of intact TPELN (parental band) of young and old VSMCs. Graph showing that individual values of relative fragment abundance and mean ± SEM. Unpaired *t* test, \*\**p* < 0.01. **B**. Representative Western blots of MFG-E8 in young and old rat VSMCs. Graph showing that individual values of MFG-E8 expression and mean ± SEM. Unpaired *t* test, \**p* < 0.04.

#### Altered MFG-E8 and Tropoelastin Expression in Aging VSMCs

In VSMCs isolated from rat arterial walls, intact TPELN protein levels significantly decreased in old VSMCs while MFG-E8 protein expression nearly tripled in VSMCs cultured from old vs. young aortae (**Figure 7A & B**).

Interestingly, treating both young and old VSMCs with a recombinant human MFG-E8 (rhMFG-E8, 100ng/ml) markedly reduced the levels of intact TPELN protein in both young and old cells (two-way ANOVA: main effect of age, *p* < 0.01; main effect of rhMFG-E8 treatment, *p* < 0.001; interaction, *p* < 0.05) (**Figure 8A**); the abundance of TPELN mRNA, however, was not significantly altered in either young or old cells (**Supplemental Figure I**). Importantly, silencing MFG-E8 mRNA markedly suppressed MFG-E8 protein levels in both young and old VSMCs (**Supplemental Figure II**), and consequently, intact TPELN protein levels were significantly increased in both young and old cells (two-way ANOVA: main effect of age, *p* < 0.0001; main effect of si-MFG-E8 treatment, *p* < 0.0001; interaction, *p* < 0.01) (**Figure 8B)**. Notably, MFG-E8 silencing did not significantly affect TPELN mRNA levels (**Supplemental Figure III**). These findings suggest that MFG-E8 associated reduction of intact TPELN is due mainly to post translational modification.

**Figure 8.**
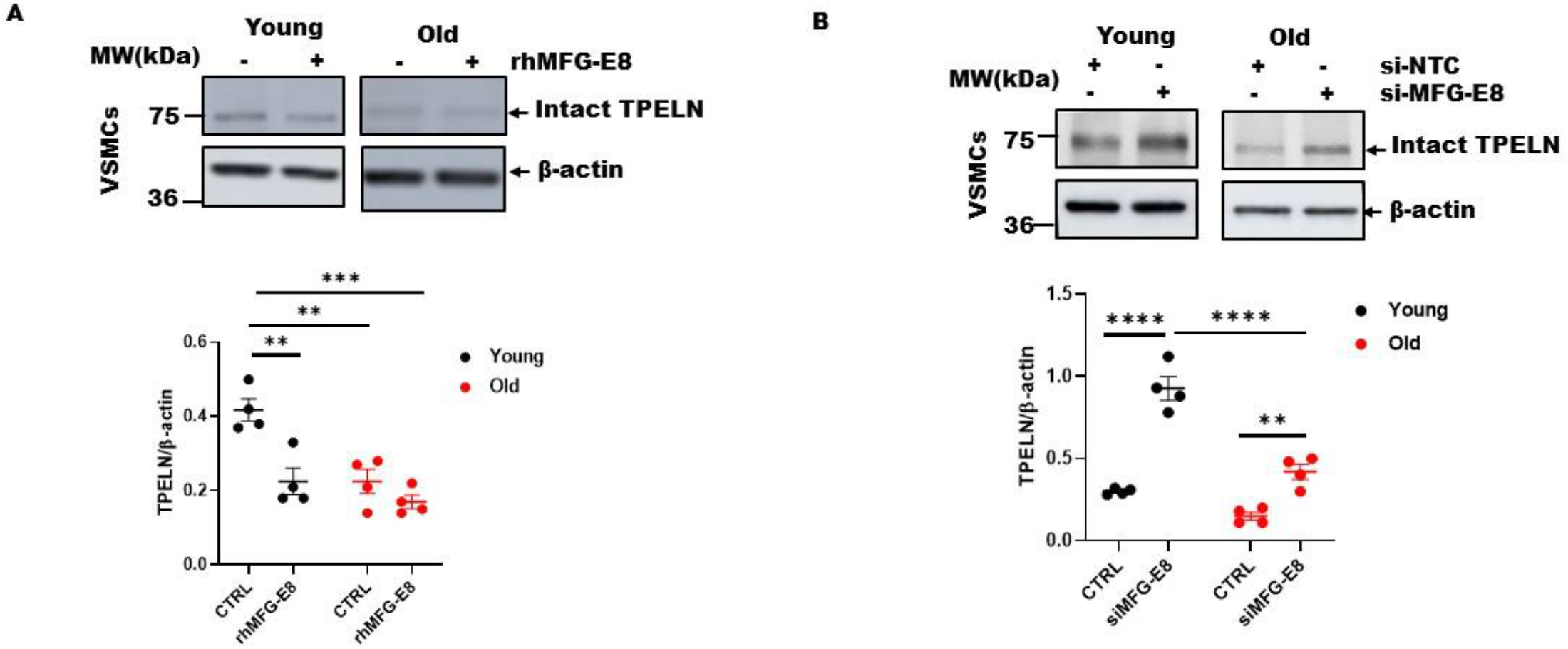
MFG-E8 reduces intact TPELN protein levels in aging rat VSMCs. **A**. Young and old VSMCs were treated for 24 hours in medium containing rhMFG-E8 (100 ng/ml). Representative immunoblots of intact TPELN in young and old VSMCs treated with rhMFG-E8 (100 ng/ml). Graph showing that individual values of TPELN expression and mean ± SEM, analyzed with two-way ANOVA (main effect of age, *p* < 0.01; main effect of genotype, *p* < 0.01, interaction, *p* < 0.05) followed by Turkey’s multiple comparison test: \*\**p* < 0.01 and \*\*\**p* < 0.001. **B**. Young and old VSMCs were transfected for 24 hours with scrambled si-RNA (20 nM) (si-NTC) or si-MFG-E8 (20 nM). Representative immunoblots of intact TPELN of homogenous lysates from young and old VSMCs treated with siRNA. Graph showing that individual values of TPELN abundance and mean ± SEM, analyzed with two-way ANOVA (*p* < 0.0001, main effect of age; *p* < 0.0001, main effect of genotype; interaction, *p* < 0.01) followed by Turkey multiple comparison test: \**p* < 0.05.

#### MFG-E8 Activates MMP-2 in Aging VSMCs

In situ gelatin zymogram demonstrated that gelatinase activity was markedly increased in old versus young VSMCs, predominantly located in the plasma membrane (**green, Figure 9A)**. PAGE zymogram further indicated that the increased activated gelatinases were activated MMP-2 (**Figure 9B**). Remarkably, an exposure of either young or old VSMCs to rhMFG-E8 for 24 hours was markedly increased MMP-2 activation (two-way ANOVA: main effect of age, *p* < 0.0001; main effect of rhMFG-E8 treatment, *p* < 0.0001; interaction, *p* < 0.01) (**Figure 9C**)

**Figure 9.**
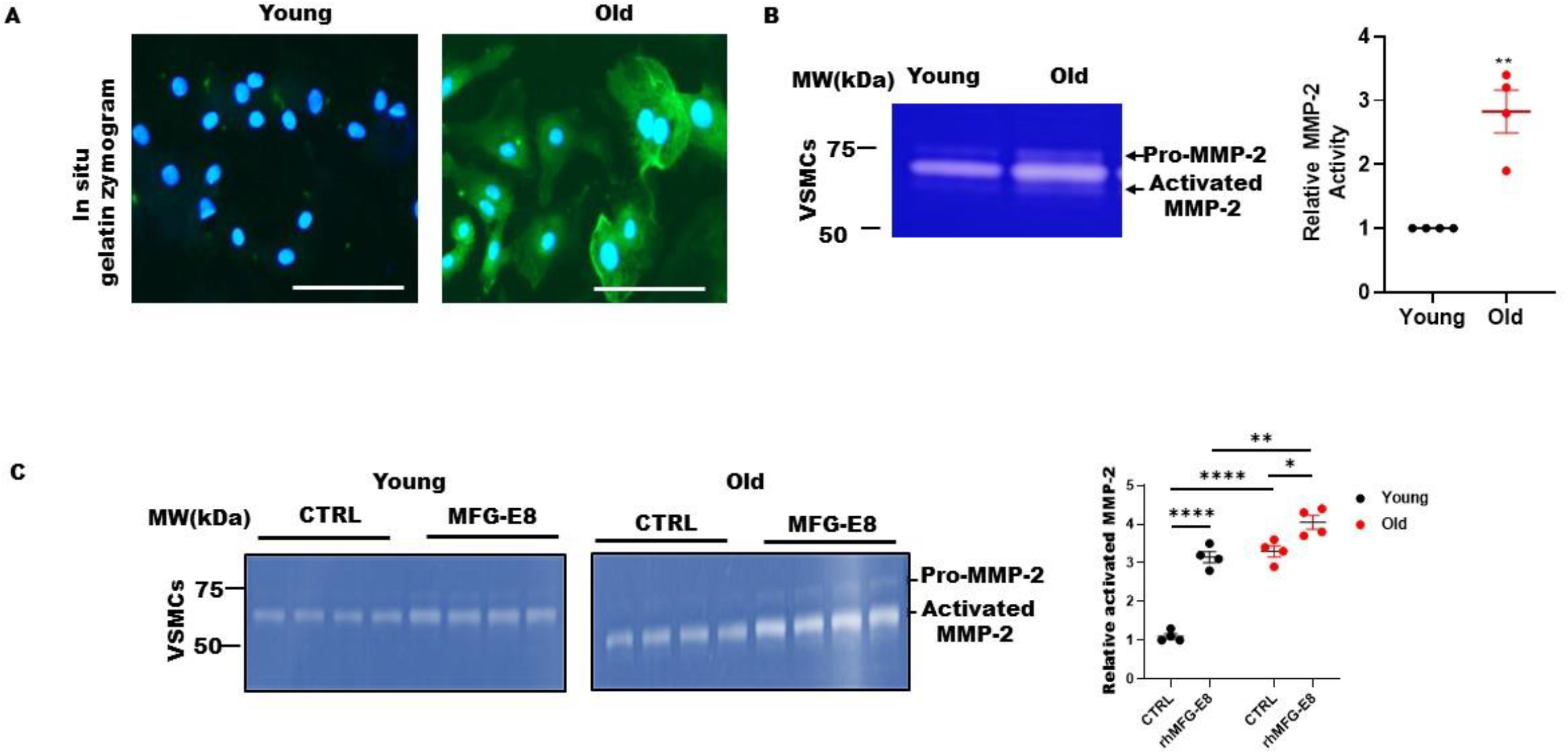
MMP-2 activation in aging rat VSMCs and MFG-E8 treatment. **A**. In situ zymogram of young and old VSMCs (green). Bar = 100μm. **B**. PAGE zymogram of young and old VSMCs. Graph showing that individual values of activated MMP-2 and mean ± SEM followed by unpaired *t* test, \*\**p* < 0.01. **C**. Representative zymogram of medium collected from cultured young and old VSMCs treated with or without rhMFG-E8 for 24 hours (100 ng/ml) (left side). Graph showing that individual values of activated MMP-2 and mean ± SEM: Graph showing that individual values of TPELN expression and mean ± SEM, analyzed with two-way ANOVA (main effect of age, *p* < 0.0001; main effect of treatment, *p* < 0.0001, interaction, *p* < 0.001) followed by Turkey’s multiple comparison test: unpaired *t* test, \**p* < 0.05.

#### MMP-2 Cleaves TPELN in Aging VSMCs

Co-immunoprecipitation revealed that MMP-2 protein physically interacted with intact TPELN in young VSMCs but was undetectable in old cells because intact TPELN was almost completely degraded by the activated MMP-2 **(Figure 10A**). Indeed, our in vitro experiments showed that exogenous activated MMP-2 effectively cleaved intact TPELN in VSMCs (**Figure 10B**). Notably, silencing MMP-2 significantly enhanced intact TPELN protein abundance in both young and old cells (two-way ANOVA: main effect of age, *p* < 0.0001; main effect of siMMP-2, *p* < 0.01; interaction of age x siMMP2, *p* > 0.01) (**Figure 10**).

**Figure 10.**
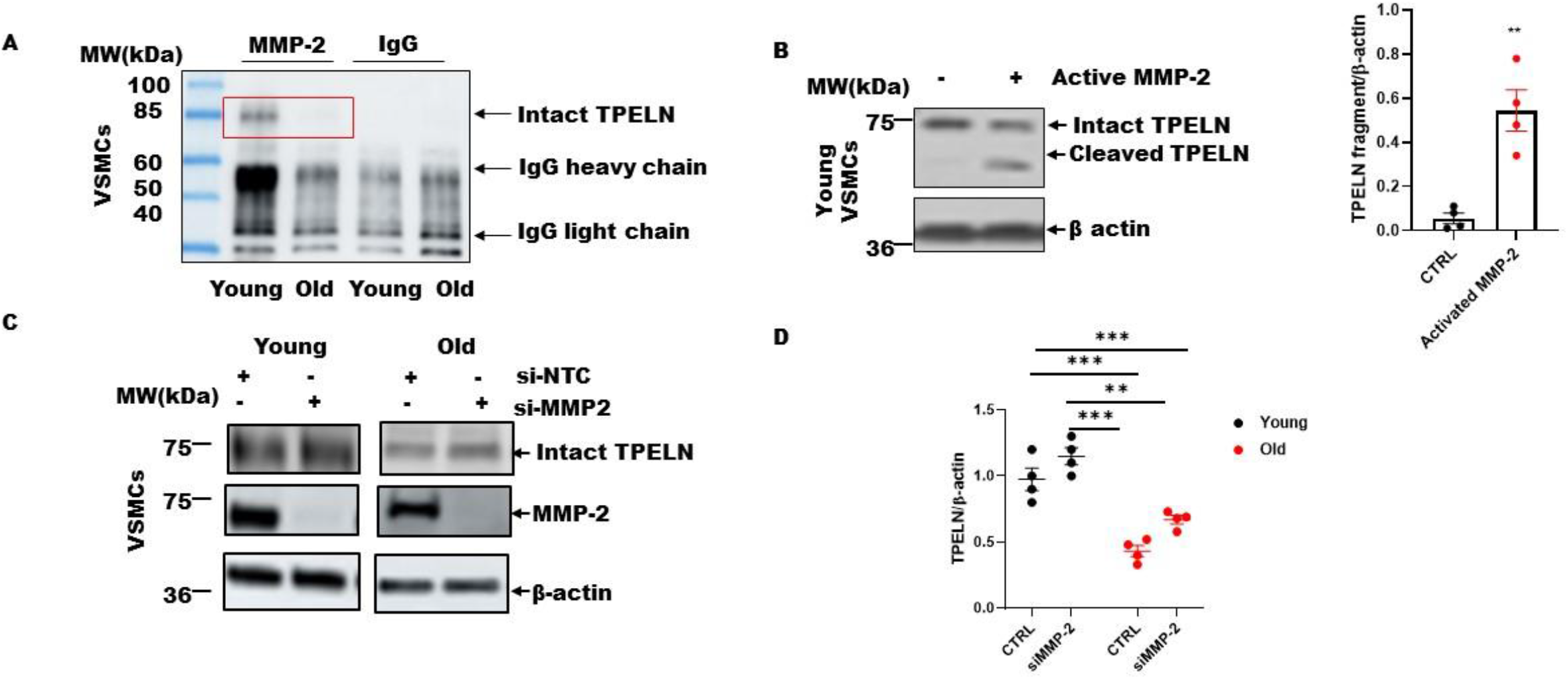
MMP-2 physically interacts with and degrades TPELN in aging rat VSMCs. **A**. Representative TPELN immunoblotting of MMP-2 immunoprecipitants from young and old VSMC cell lysates. **B**. Representative immunoblotting (left panel) of young VSMC cell lysates. Cells were treated with activated MMP-2 for 3 hours at 37°C. Graph showing that individual values of TPELN fragment and mean ± SEM. Unpaired *t* test, \**p* < 0.05. **C**. Representative immunoblotting of TPELN and MMP-2 from young and old VSMC cell lysates treated with si-NTC (20 nM) or siMMP-2 (20nM). **D**. Graph showing that individual values of TPELN abundance and mean ± SEM, analyzed with two-way ANOVA (*p* < 0.0001, main effect of age; *p* < 0.01, main effect of treatment; *p* > 0.05, interaction) followed by Turkey’s multiple comparison test: \*\**p* < 0.01.

## Discussion

MFG-E8 plays an important role in arterial aging.^20–23,32^ The current findings demonstrated that MFG-E8 actively participates in age-associated structural and functional aortic remodeling, including elastic fiber degeneration, collagen deposition, and calcification in vivo; and MFG-E8 activates MMP-2, and subsequently degrades TPELN secreted from VSMCs in vitro. We also found that the MFG-E8 deficiency alleviates aortic remodeling through an inhibition of elastic fiber degradation and calcification. These findings reveal that MFG-E8 signaling promotes age-associated arterial elastolysis, fibrosis, and calcification.

The age-associated increase in MFG-E8 facilitates elastic fiber degradation. Age increases MFG-E8, accompanied by an elevation in elastin degradation within the aortic wall in rats and mice. However, the absence of MFG-E8 alleviates the age-associated increase in elastic fiber degradation. The current in vitro findings indicated that 1) MFG-E8 treatment significantly impacted on TPELN protein level but didn’t alter TPELN mRNA levels of VSMCs; 2) MFG-E8 treatment significantly decreased intact TPELN levels, which was accompanied by an increase in activated MMP-2; and 3) Activated MMP-2 effectively cleaved intact TPELN in VSMCs; in contrast, silencing MMP-2 increased intact TPELN levels in VSMCs. TPELN is a monomeric soluble precursor of insoluble elastin secreted from VSMCs, which is located to on the surface of plasma membrane and assembled into a mature elastic fiber, maintaining elastic network integrity.^14^ These findings suggest that MFG-E8 promotes elastic fiber degradation in the arterial all with aging. The age-associated increase in MFG-E8 abundance promotes calcification in the arterial wall. Aging increases arterial calcification in rats and mice, accompanied by increases in fibrosis and elastolysis. The absence of MFG-E8 significantly inhibited fibrosis and elastin degradation, and importantly mitigated arterial calcification in mice with aging.

Aging significantly increases MMP-2 activity, TGF-β1 activation, p-SMAD-2/3, collagen type I and type II deposition, elastic fiber degradation in the arterial wall in rats.^21,26,33^ Deposited collagen (types I & II) and degraded elastic fibers are major substrates of calcification in the arterial wall.^14,34^ The fractured ends or eroded points of degraded arterial elastin fragments have a high affinity for calcium-phosphorus products, which become a nucleation site for the deposition of hydroxyapatite crystals, an impetus for propagating arterial calcification.^8,9,12,31^ Indeed, the fracture of elastic fibers and TPELN cleavage is closely associated with calcification both in vitro though VSMCs and *in vivo* in the arterial wall.^10,22,35,36^ We also found that levels of both ALP and Runx2 were markedly reduced in old KO versus WT. ALP, a non-tissue specific enzyme, is a necessary biomineralizing molecule, and facilitates calcification in the arterial wall.^11,30,37^ Runx2, a transcription factor, enhances the chondro-osseous differentiation of VSMCs and promotes calcification and stiffness in the arterial wall.^38^ These findings reveal that MFG-E8 signaling modifies arterial calcification with aging.

In summary, the age-associated increase in MFG-E8 protein abundance promotes adverse arterial wall remodeling, including elastic fiber degradation, collagen deposition, calcification in the arterial wall with aging. MFG-E8 associated increases in activated MMP-2, ALP, and RUNX2 in VSMCs are potential cellular and molecular events promoting elastolysis, fibrosis and calcification in the aging arterial wall. Thus, targeting MFG-E8 molecule is a novel molecular therapeutic approach to delaying arterial aging, maintaining arterial health.

## Acknowledgments

The authors thank Kimberly Raginski and Robert Monticone for editing this manuscript and Cristopher Morrel for assisting with statistical analysis. Additionally, the authors are grateful to Yolanda L. Jones, NIH Library, for additional editing.

## CLINICAL PERSECTIVE

### What Is New?

- Milk fat globule epidermal growth factor VIII (MFG-E8) is a proinflammatory molecule in the arterial wall with aging.
- MFG-E8 is closely associated with age-associated extracellular matrix remodeling in the arterial wall.
- MFG-E8 deficiency reduces age-associated arterial elastolysis, fibrosis, and calcification.

### What Are the Clinical Implications?

The inhibition of MFG-E8 is a novel therapeutic approach to the prevention or treatment of elastolysis, fibrosis, and calcification in age-associated arterial remodeling.

