## Supplementary material for "Effects of Milk Fat Globule Epidermal Growth Factor VIII On Age-Associated Arterial Elastolysis, Fibrosis, and Calcification": Suppl Materials and Figure

**Running title:** MFG-E8 is Essential for Arterial Aging

*Equal contribution

Correspondence to Mingyi Wang, MD, PhD, FAHA

Laboratory of Cardiovascular Science

National Institute on Aging

National Institutes of Health

Biomedical Research Center (BRC)

251 Bayview Boulevard, Baltimore, MD 21224, USA

Tel: (+1)410-558-8112

**Keyword:** Aging ● Vascular smooth muscle cells ● Milk fat globule epidermal growth factor VIII ● Matrix metalloprotein type II ● Tropoelastin ●Elastolysis ● Calcification● Fibrosis

**Subject codes:** Vascular biology● Inflammation● extracellular remodeling

**Supplemental Figure Legends**

**Supplemental Figure I. Effects of rhMFG-E8 on TPELN transcription.** TPELN mRNA levels from both young and old rat VSMCs as determined by RT-PCR. Graphs shows individual TPELN mRNA values and mean ± SEM. Unpaired *t* test: not significant.

**Supplemental Figure II. MFG-E8 silencing in aging VSMCs. A.** MFG-E8 mRNA levels (left panel) as determined by RT-PCR on both young and old rat VSMCs treated with both si-MFG-E8 and si-NTC control. Graphs shows individual TPELN mRNA values and mean ± SEM. Unpaired *t* test: ***$p < 0.01$. **B.** Immunoblotting of MFG-E8 in both young and old VSMCs treated with both si-MFG-E8 and si-NTC control.

**Supplemental Figure III. Effects of si-MFG-E8 on TPELN transcription.** TPELN mRNA levels from both young and old rat VSMCs treated with both si-MFG-E8 and si-NTC control as determined by RT-PCR. Graphs shows individual TPELN mRNA values and mean ± SEM. Unpaired *t* test: not significant.

RT-PCR

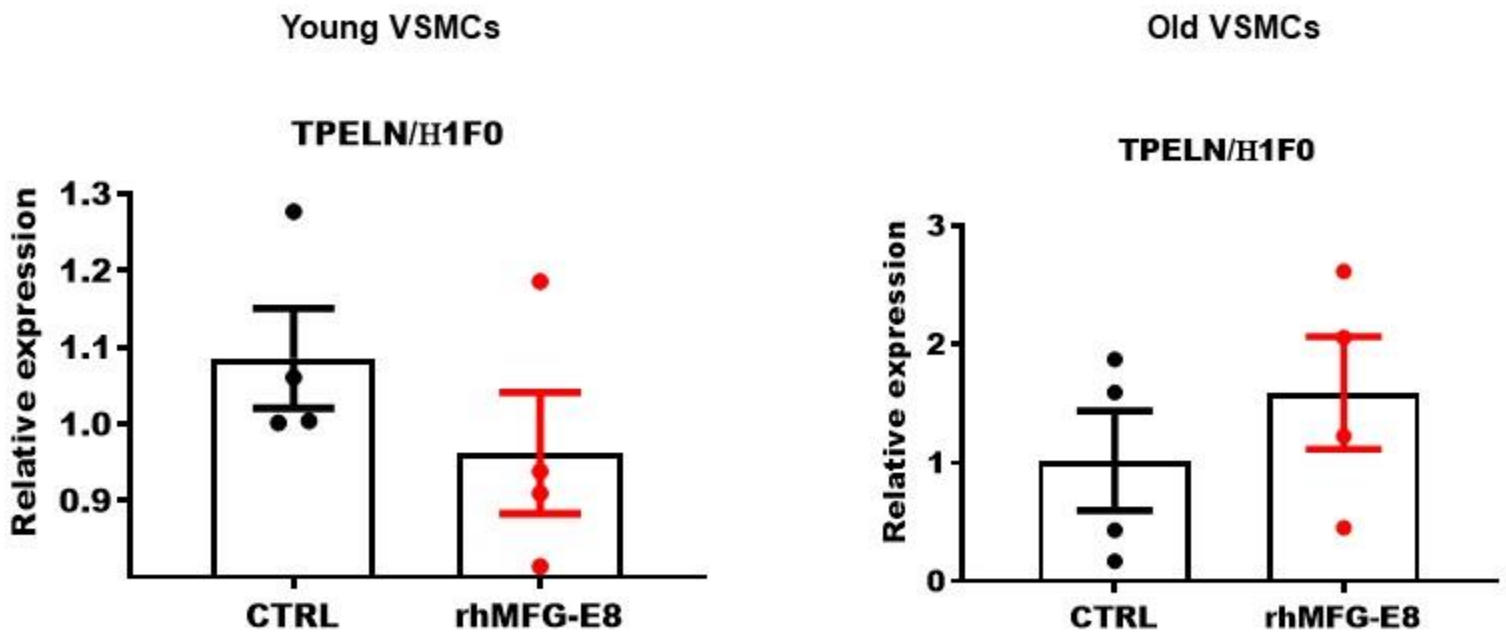


Supplemental Figure I

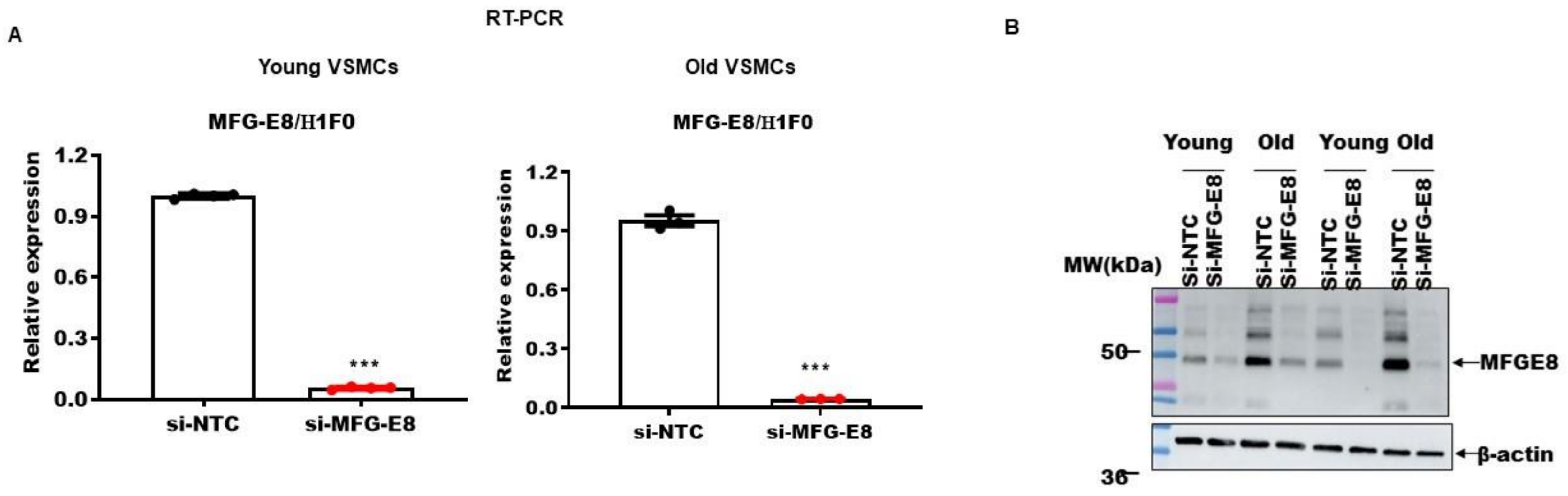


Supplemental Figure II

RT-PCR

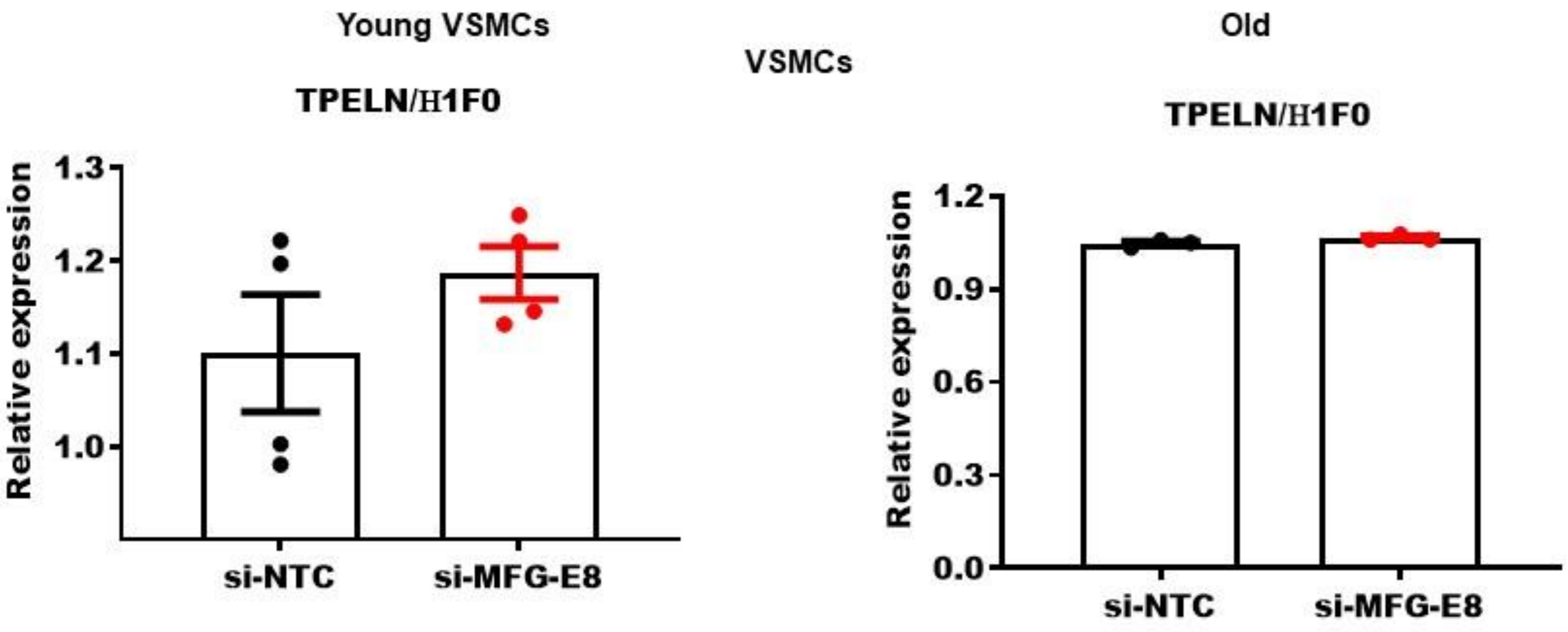


Supplemental Figure III
